# Expression of a gene for an MLX56 defense protein derived from mulberry latex confers strong resistance against a broad range of insect pests on transgenic tomato lines

**DOI:** 10.1101/2020.09.17.301143

**Authors:** Mika Murata, Kotaro Konno, Naoya Wasano, Atsushi Mochizuki, Ichiro Mitsuhara

**Affiliations:** Institute of Vegetable and Floriculture Science, National Agriculture and Food Research Organization (NARO), Mie Prefecture, Japan; Institute of Agrobiological Sciences, NARO, Ibaraki Prefecture, Japan; Department of Science, University of Toyama, Toyama Prefecture, Japan; Institute of Agro-Environmental Sciences, NARO, Ibaraki Prefecture, Japan

## Abstract

Insect pests cause serious damage in crop production, and various attempts have been made to produce insect-resistance crops, including the expression of genes for proteins with anti-herbivory activity, such as BT toxins. However, the number of available genes with sufficient anti-herbivory activity is limited. MLX56 is an anti-herbivory protein isolated from the latex of mulberry plants, and has been shown to have a strong growth-suppressing activity against the larvae of a variety of lepidopteran species. As a model of herbivore-resistant plants, we produced transgenic tomato lines expressing the gene for MLX56. The transgenic tomato lines showed strong anti-herbivory activities against the larvae of the common cutworm, *Spodoptera litura*. Surprisingly, the transgenic tomato lines also exhibited strong activity against the attack of the western flower thrips, *Frankliniera occidentalis*. Further, growth of the hadda beetle, *Henosepilachna vigintioctopunctata* fed on leaves of transgenic tomato was significantly retarded. The levels of damage caused by both western flower thrips and hadda beetles were negligible in the high-MLX56-expressing tomato line. These results indicate that introduction of the gene for MLX56 into crops can enhance crop resistance against a wide range of pest insects, and that MLX56 can be utilized in developing pest-resistance GM crops.

## Introduction

Damage due to feeding insects has a large impact on crop production, reducing both the amount and quality of the product. Also, herbivore damage enhances vulnerability to infection, and some herbivores act as vectors of pathogens. Therefore, considerable costs have been expended to protect crops from herbivores, and protective agents against herbivores are continuously being developed. The primary protective agents against herbivores have been pesticides, and some insect-resistant trails have been introduced by traditional breeding. In recent decades, many transgenic approaches have been taken to produce herbivore-resistant crops by introducing genes for anti-insect proteins, such as inhibitors of the digestive enzymes of herbivores or lectins. Most attempts, however, have been unsuccessful in practical use, except for the cases of Bt (*Bacillus thuringensis*) toxins, which showed sufficient toxicity to pests in very low concentrations [1].

Introduction of genes for Bt toxins such as Cry1Ab, resulted in strong resistance against insect pests sensitive to Bt toxins. Many transgenic crops into which the genes for Bt toxins were introduced, such as maize, soybean, and cotton, are commercially successful and cultivated worldwide [1]. However, each of the Bt toxins is effective only for a limited range of insect pests, such as the corn earworm, *Helicoverpa zea* (Lepidoptera: Noctuidae), and the European corn borer, *Ostrinia nubilalis* (Lepidoptera: Crambidae) [2]. Indeed, no Bt proteins are known to be effective against all lepidopteran insects. To obtain transgenic crops with resistance against various types of insect pests at the same time, it would be necessary to introduce multiple Bt genes (proteins) with different toxic spectra simultaneously. In addition to the drawback that Bt has a narrow toxic spectrum against insect pests, the incidence of Bt-resistant pests is increasing [3, 4]. Therefore, there is a great need for the discovery of novel anti-insect proteins with strong anti-insect activities and broad toxic spectrum against insect pests.

The MLX56 protein was isolated from the latex of mulberry trees as a potent growth-suppressing protein active against the larvae of lepidopteran species [5]. MLX56 is a highly glycosylated protein with an apparent molecular mass of around 56kDa. It has a unique structure compared to other anti-insect proteins with an extensin domain, which is proline-rich and highly arabinosylated, surrounded by two chitin-binding domains (hevein domains) in its N’ region, and in C’ region, with a chitinase-like domain containing mutations that result in an absence of chitinase activity. MLX56 inhibited the growth of the larvae of lepidopteran species such as the cabbage armyworm, *Mamestra brassicae* (Lepodoptera: Noctuidae) and the Eri silkworm, *Samia ricini* (Lepodoptera: Satuniidae) at extremely low concentrations (0.01-0.03% / wet artificial diet) [5]. In addition, MLX56 and its close homolog LA-b showed toxicity to the larvae of fruit flies reared on a diet containing MLX56 [6, 7]. These observations suggest that the MLX56 family proteins (MLX56 and its homologs) have strong anti-herbivore activity against a wide range of insects. Interestingly, growth of the silkworm, *Bombyx mori*, a mulberry specialist, was not at all suppressed by MLX56 or by MLX56-containing mulberry latex, suggesting that as a mulberry specialist *B. mori* may have developed some adaptive mechanism to MLX56 and other latex toxins (e.g., sugar-mimic alkaloids) as mulberry specialist [5, 8, 9, 10].

A recent study showed that not only the MLX56 structure, but also the mode of the anti-insect action of MLX56 is unique [11]. Specifically, the study showed that the peritrophic membrane (PM; a thin membrane wrapping food material in the midgut of insects) in the midgut of the *Eri silkworm, S. ricini*, exhibited abnormal swelling when fed a diet containing MLX56. The findings suggested that the hevein domains of MLX56 bind to chitin containing PM, that the swelling of PM is induced by the swelling activity of the extensin domain, and that the swollen thick PM suppresses insect growth by functioning as a barrier against the movement of nutrient and digestive enzymes in the midgut of insects [11, 12]. MLX56 may inhibit the growth of insects belonging to taxa other than Lepidopteran in similar ways, because insects belonging to most insect taxa have PM in their midgut. However, insects belonging to taxa such as Thysanoptera lack PM [13], and therefore whether or not MLX56 can exert a growth inhibitory activity against thrips is an interesting open question.

Thrips attack a wide range of crops worldwide and cause serious damage to production, especially in greenhouse cultivation. Control of the thrips is difficult because of their tiny body sizes and ability to develop pesticide resistance [14, 15]. In addition, thrips act as vectors of viral diseases. For instance, the western flower thrips, *Frankliniera occidentalis* (Thysanoptera: Thripidae), is known to be a vector for tospoviruses, including tomato spotted wilt virus (TSWV) and impatiens necrotic spot virus (INSV) [16, 17]. Therefore, novel protection methods to control thrips are needed, and it seems worthwhile to examine whether MLX56 could be effective. Meanwhile, the hadda beetle, *Henosepilachna vigintioctopunctata* (Coleoptera: Henosepilachna), is known to be an important pest in Solanaceae crops such as potato and eggplant in several countries, including Taiwan, China and India [18, 19]. Screening of varieties for resistance to *H. vigintioctopunctata* has been studied [20, 21]. Hence, a breakthrough in control measures against this pest is also urgently needed.

Since the growth-inhibitory activity of MLX56 against insect herbivores is evident even at very low concentrations (ca. 0.01%-0.03%), MLX56 appears to be a useful protein in the effort to produce GM plants resistant to insect pests. We previously showed that transient expression of the gene for MLX56 can enhance plant resistance against the common cutworm, *Spodoptera litura* (Lepidoptera: Noctuidae), the cabbage armyworm, *M. brassicae*, and the diamondback moth, *Plutella xylostella* (Lepidoptera: Plutellidae) in tobacco, tomato or *Arabidopsis* plants [22]. In the present study, we produced transgenic tomato lines expressing the gene for MLX56, and then tested for resistance against the common cutworm (Lepidoptera), the western flower thrips (Thysanoptera) and the hadda beetle (Coleoptera).

## Methods

### Production of a transgenic tomato plant

A binary vector with a strong constitutive promoter::MLX56 gene [22] was used for the tomato plant transformation. The plasmid was introduced into Agrobacterium LBA4404 as an intermediate host. *Agrobacterium*-mediated transformation of the tomato plant cv. Micro-Tom was performed as described in Sun et al. (2006) [23]. A transgenic tomato plant transformed with the same promoter::Luciferase construct was also produced as a vector control plant.

Regenerated plants were planted in pots and grown in a growth chamber controlled at 25 °C and a 16h /8h light/dark photoperiod. Expression of the transgene was determined by qRT-PCR as described by Kawazu et al. (2012) [22]. Transgenic tomato plants with high-level expression of the gene for MLX56 were selected and transferred to an isolated greenhouse. Seeds obtained from primary transgenic plants were seeded onto kanamycin containing plates and kanamycin resistant progenies were used for further analysis.

### Partial purification and detection of MLX56 protein

Leaves of second-generation of MLX56 expression tomato plants and vector control plant were excised, and 200 mg of fresh leaf pieces were homogenized with 800 μL of 20 mM Tris-HCL (pH 9.5) supplemented with protease inhibitor cocktail (Complete Mini, Roche) using a mortar and pestle. The homogenate was centrifuged at 12,000 rpm for 5 min, and the supernatant was obtained as a “crude extract”.

A slurry of chitin beads (New England BioLabs) preequilibrated to and then suspended in 2 vol. of the extraction buffer, and then 200 μL of a slurry of chitin beads and 200 μL of crude extract were each mixed and incubated for 10 min at room temperature. Then the mixtures were centrifuged at 12,000 rpm for 5 min to separate “unbound sap” and protein-bound chitin beads. The chitin beads were washed 3 times with extraction buffer and the wash solution was deposited as the “wash fraction”. The chitin beads were then further washed with 200 μL of 8 M of urea solution, and the urea supernatant was deposited as the “urea fraction”. Proteins tightly bound on the chitin beads were eluted and denatured by resuspension in 200μL of SDS sample buffer and by boiling for 5 min. Crude extract and the above fractions were then denatured in SDS-PAGE buffer for SDS-PAGE separation. Samples were separated by SDS-PAGE using a Mini-Protein Tera Cell system (Bio-Rad Laboratories, Inc., Hercules, USA, CA). Proteins were visualized by CBB staining as described elsewhere.

### Insect resistance assay

Eggs of cotton cutworm (*S. litura*) were purchased from Sumika Techno Service Co. (Takarazuka, Japan). Tomato leaves were excised with scissors, and two leaves were put in a 1.5mL microtube filled with distilled water to prevent leaves from drying. Ten hatchlings of *S. litura* were inoculated onto the two tomato leaves in a plastic container (15cm diameter × 8cm high) at 25 ± 1 °C with a 14L/10D photoperiod. The numbers of surviving larvae and instar numbers were recorded after 3, 6 and 9 days. Differences in the numbers of surviving larvae and the proportion of the instar numbers were statistically compared using Tukey-Kramer HSD test after one-way ANOVA, and Ryan’s multiple-range test for proportions after the *χ*^2^ test, respectively.

For evaluation of tomato resistance against the western flower thrips (*F. occidentalis*), 3-4 week-old potted tomato plants were transferred into separate transparent vessels (bottom diameter 8.5 cm, top diameter 10.5 cm, height 14 cm) separately. Twenty adult female thrips were placed in each vessel and capped. The cap of the vessel had a window covered with nylon mesh for ventilation. The vessels containing tomato were incubated in a growth chamber at 25 ± 1 °C with a 14L/10D photoperiod. The numbers of surviving adults, pupae and larvae were determined after two weeks. Fifteen plants were used for each strain of transgenic tomato line. Data on survival rates were analyzed by Tukey-Kramer HSD test after one-way ANOVA.

Hadda beeltles (*H. vigintioctopunctata*) that were collected in Tsukuba, Japan, and then kept in the laboratory in Tsukuba for two generations were used for bioassays. Each group of 12 newly hatched first instar larvae was fed excised leaves of either the control line (Micro-Tom) or the transgenic lines (line 69 or line 73) for 9 days in an incubator at 25 °C with a 16L/8D photoperiod. Leaves were replaced by new ones every other day, the body weight of each larvae was measured, and then the data were statistically analyzed using Tukey’s test for multiple comparisons.

These analyses were conducted using R version 4.0.2 [24].

## Results

### Production of the MLX56-overproducing tomato plant

A tomato plant (cv. Micro-Tom) was transformed with MLX56 containing the expression vector pE12Omega. The expression vector enables a high expression of foreign gene monocot and dicot plants [25, 26]. Expression of the gene for MLX56 in each regenerated tomato plant was analyzed by qRT-PCR, and two lines, MLX56-69 and −73, were selected as highly expressing strains. Second-generation seedlings of each transgenic line were selected by kanamaycin, and levels of mRNA for MLX56 were determined at the 4 leaves stage (Fig. 1). High-level expression of the transgene was confirmed, and the levels of the mRNA for in MLX56-73 was approximately double that in MLX56-69.

**Fig. 1.**
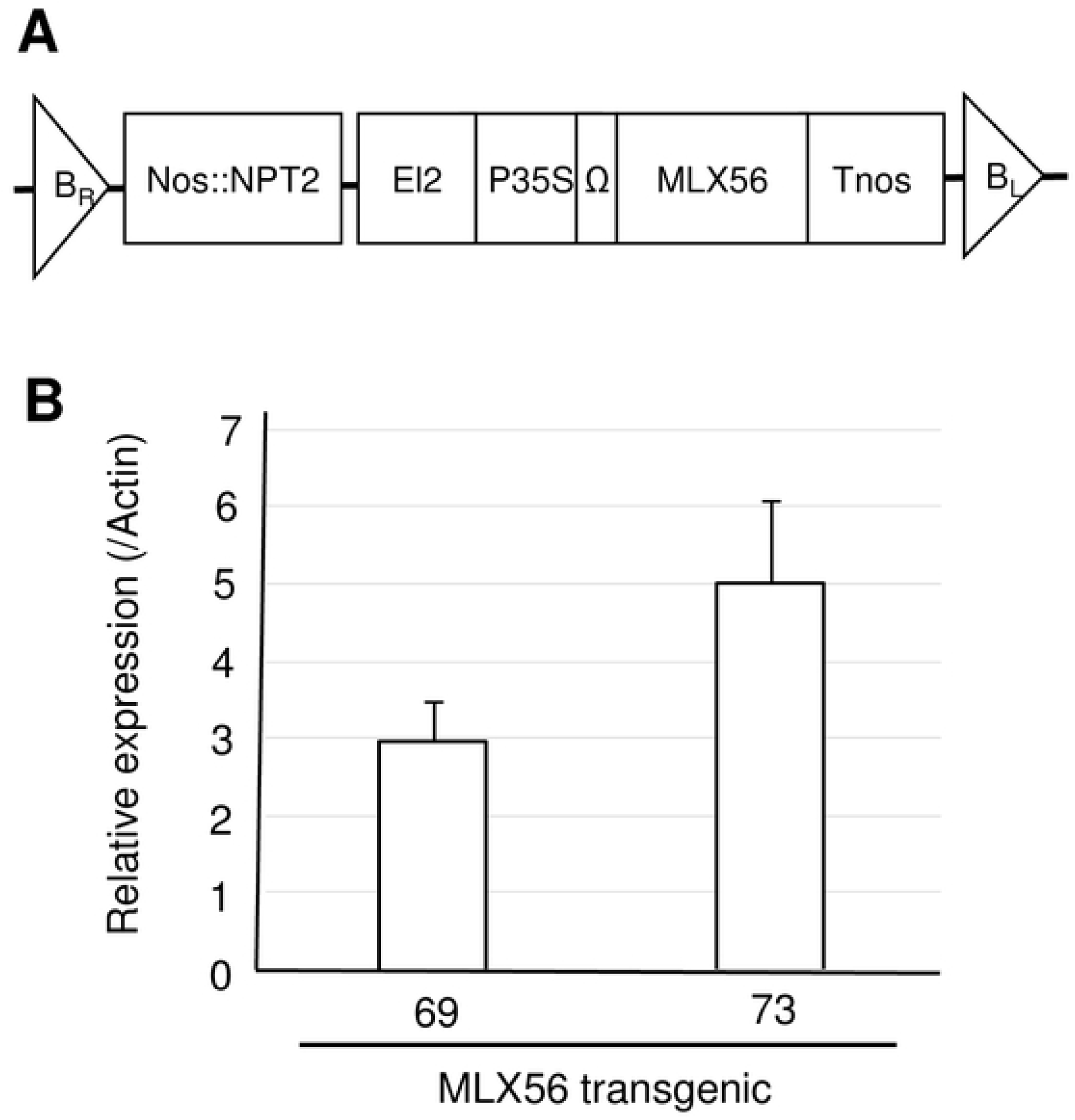
Production of the transgenic tomato plant. Structure of the introduced transgene: the expression vector pEl2Omega with the gene for MLX56 (Kawazu et al., 2012) was used for tomato transformation. The vector has the duplicated CaMV 35S transcriptional enhancer and TMV omega translational enhancer for overproduction of the transgene (A). Detection of mRNA for MLX56 in transgenic tomato leaves: the level of mRNA for MLX56 was determined by qRT-PCR using the primers, MLX56RT51 (5’-CCAAGTCCACCTCCACCAAGTC-3’) and MLX56RT31 (5’-TTTCCGAGGGCTCTTCCACATC-3) (Kawazu et al., 2012). The level of mRNA for actin was used as an internal control (B).

The growth of each transgenic line was compared to that of the control plants with the pE12Omega-containing gene for luciferase. Total weights of the transgenic lines grown in a growth chamber were not different from those of controls. Further, the yields of fruits harvested in the greenhouse were also similar between the transgenic lines and the controls.

### Detection of MLX56 protein in the transgenic tomato plant

Unfortunately, we could not obtain antibodies against the MLX56 protein with enough specific activity, and so we could not confirm production of the MLX56 protein by gel blot analysis. Instead, we tried to detect MLX56 protein as a chitin-binding protein. Chitin-binding proteins in the leaf extract of transgenic plants with the MLX56 gene or vector control were partially purified using chitin beads and separated by SDS-PAGE (Fig 2). A chitin-binding protein of around 56 kDa was detected from transgenic plants with MLX56 but not from control plants, suggesting that the gene for MLX56 produced a 56 kDa chitin-binding protein.

**Fig. 2.**
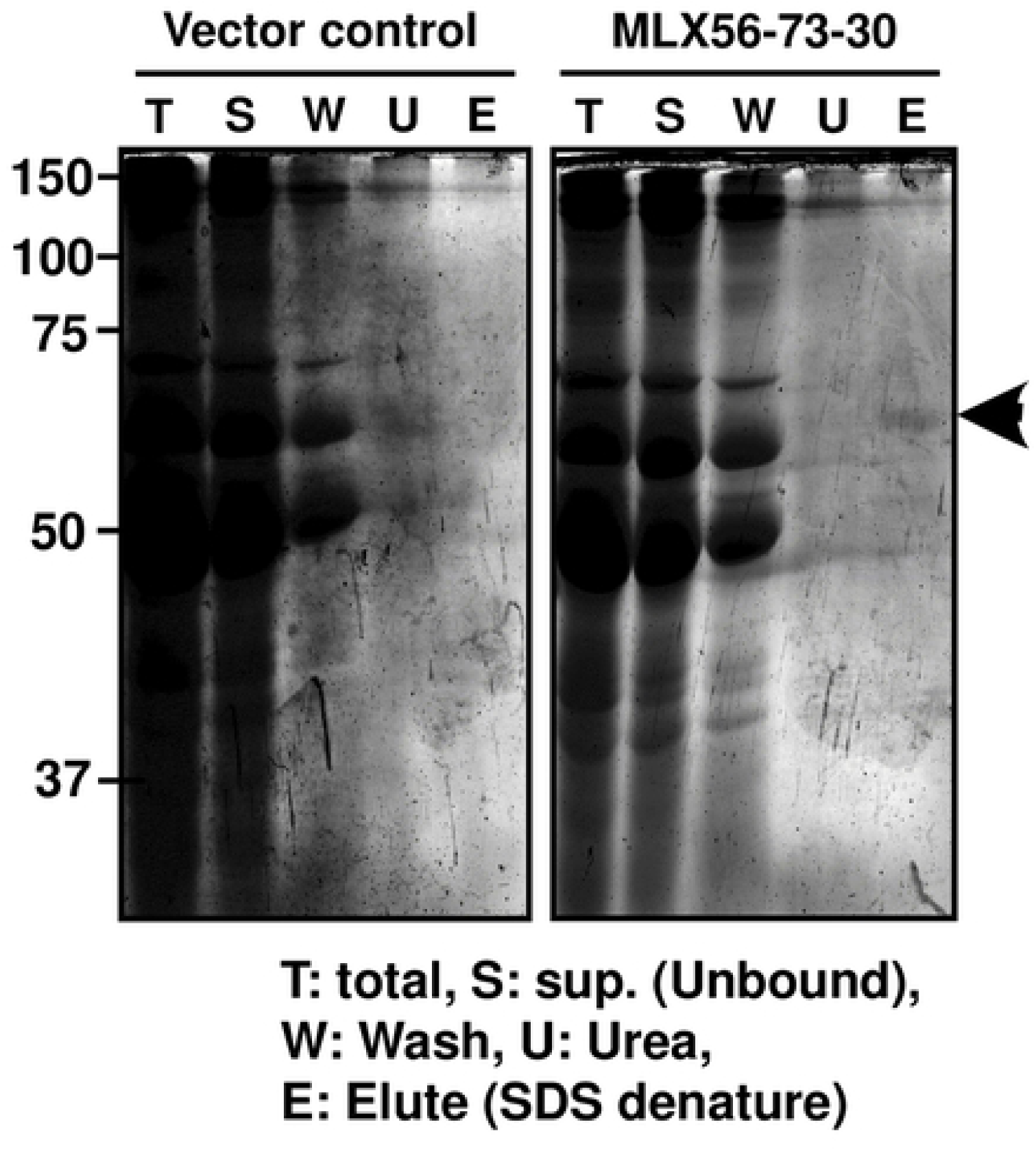
Partial purification of the chitin-binding protein in a transgenic tomato plant with the gene for MLX56. The chitin-binding protein of the control and MLX56-transformed tomato plant (MLX56-73-30) were partially purified using chitin beads. The crude extract and each partially purified fraction were separated by SDS-PAGE and visualized by CBB. T: total crude extract; S: unbound supernatant; W: wash fraction by extraction buffer; U: wash fraction by urea, E: eluted fraction after SDS denaturation that tightly bound to chitin.

### Insect resistance assay

To assess whether the MLX56-expressing tomato plants exhibited enhanced resistance against lepidopteran herbivores, larvae of *S. litura* were fed on excised leaves from each plant. The number of surviving *S. litura* larvae fed on the 69 line was significantly lower than the number of surviving larvae fed on leaves of the control on day 3, and the number of surviving larvae fed on the 73 line was significantly lower than that of surviving larvae fed on the control on days 6 and 9 (Fig. 3A) (day 3: p < 0.005, F = 8.1033, day 6: p < 0.05, F = 4.1923, day 9: p < 0.01, F = 5.8973). This confirmed the enhanced resistance against the lepidopteran herbivores by expression of the MLX56 gene. The proportion of larval instars on day 3 was significantly different from that in lines 69 and 73 (χ^2^ = 22.12, p < 0.001). On day 6, the proportion of 3^rd^ instar larvae in the control was significantly lower than that in line 69, and followed by line 73 (χ^2^ = 13.60, p < 0.005). On the other hand, there was no significant difference in the proportion on day 9 among the three lines (χ^2^ = 1.95, p > 0.05).

**Fig. 3.**
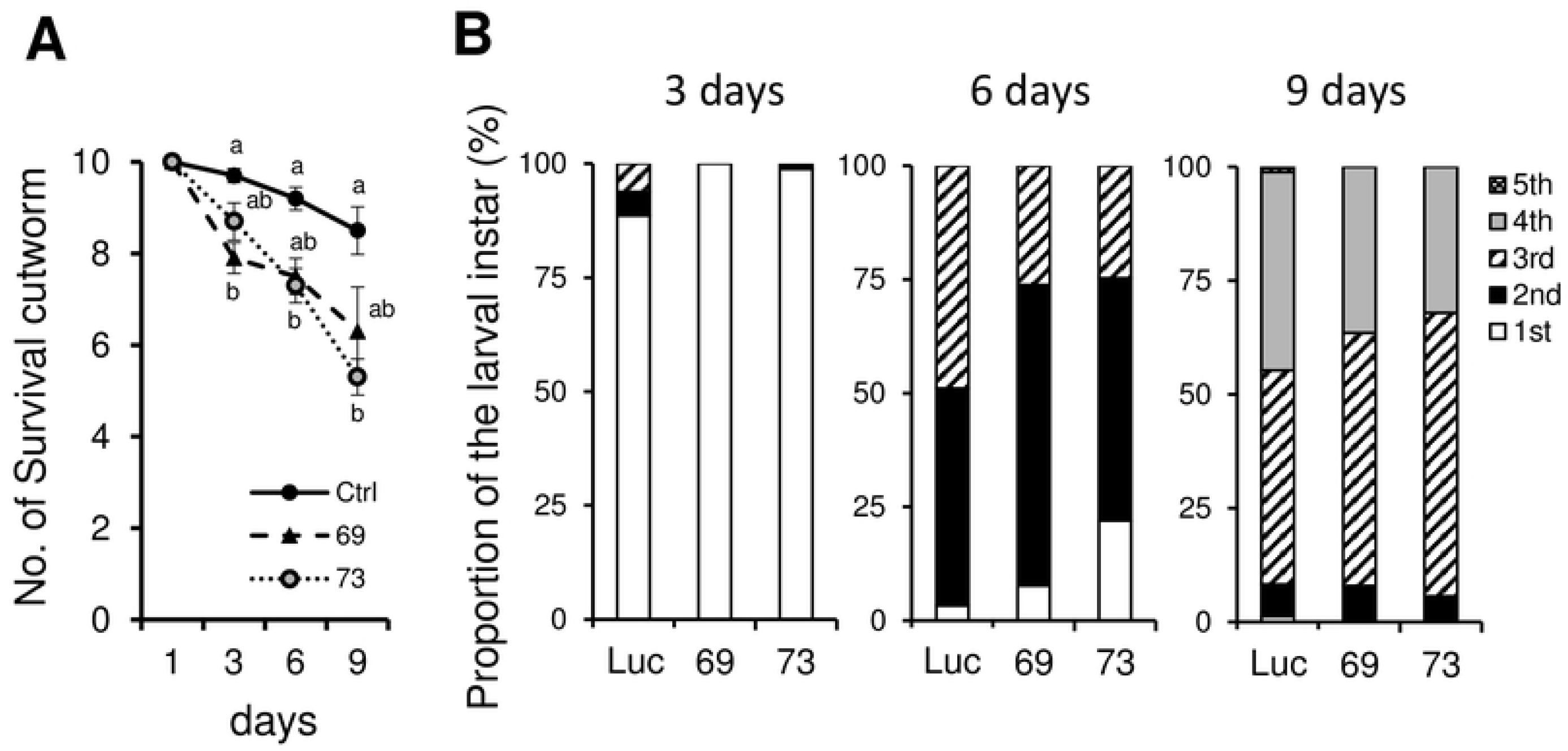
Effects of transgenic tomato on the numbers of surviving individuals and development of *S. litura* larvae. Numbers of surviving individuals from inoculation to 9 days (A). Proportions of the larval instars reared on each tomato line after 3, 6 and 9 days (B). Error bars indicate SE (n = 10). Values not followed by the same letters were significantly different among the three lines at the same day by Tukey-Kramer HSD test (day 3: p < 0.005, F = 8.1033; day 6: p < 0.05, F = 4.1923; day 9: p < 0.01, F = 5.8973).

Transgenic plants were then infested by thrips to evaluate the effect of MLX56 on herbivores other than Lepidoptera. The numbers of surviving thrips were adversely affected by transgenic tomato plants; after two weeks low numbers of surviving individuals were observed for line 73, followed by line 69 and the control (Fig. 4A) (p < 0.0001, F = 14.325). The feeding damage to tomato leaves by thrips was also less severe in line 73, followed by line 69 and the control (Fig. 4B and C). These results indicate that the expression of the gene for MLX56 is also effective against thrips.

**Fig. 4.**
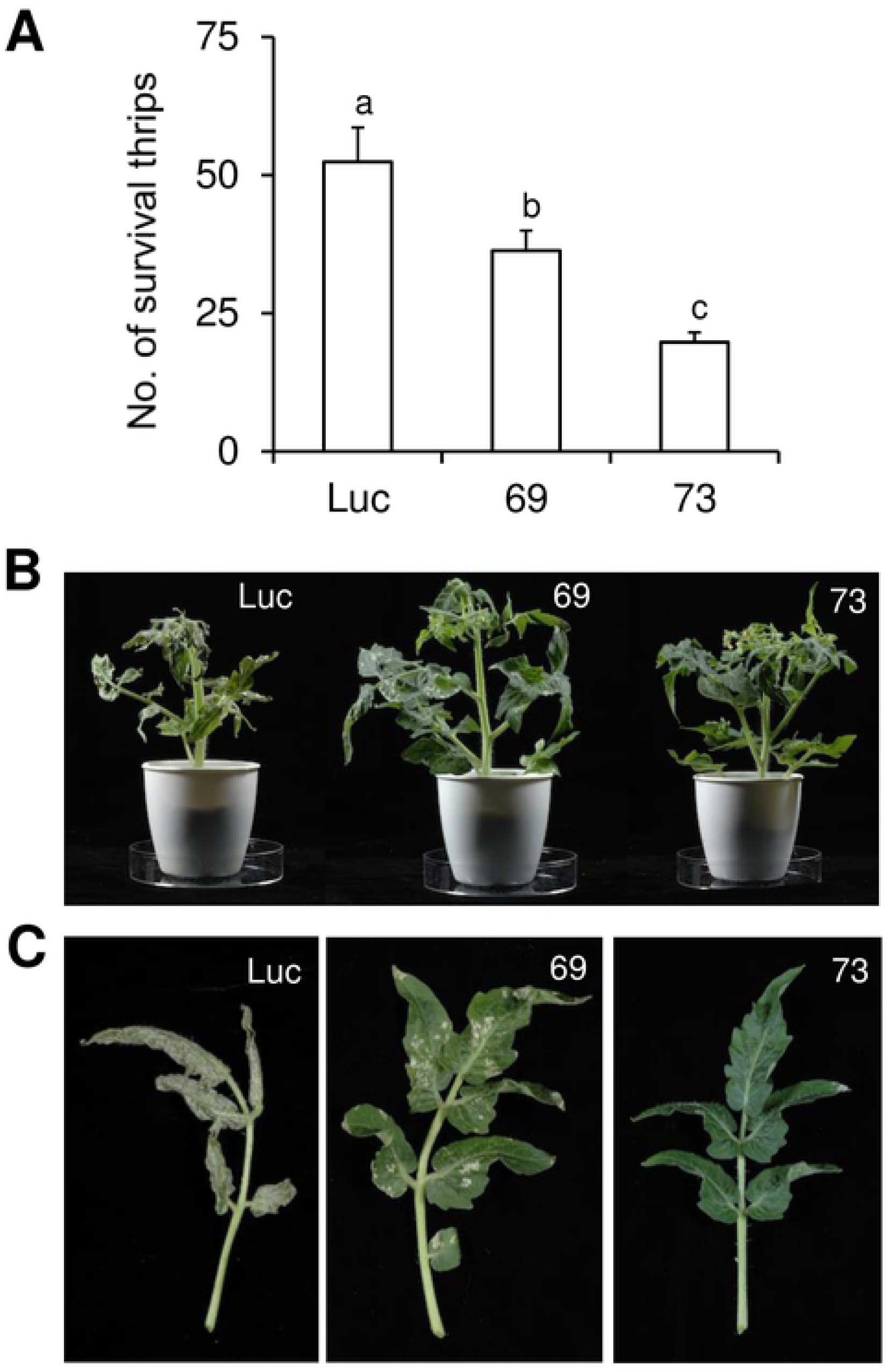
Effects of transgenic tomato on the numbers of surviving individuals of *F. occidentalis* (A), and the damage to tomato plants (B) and leaves (C) by the thrips after two weeks. Error bars indicate SE (n = 15). Values not followed by the same letters were significantly different (Tukey-Kramer HSD test: p < 0.0001, F = 14.325).

Regarding the hadda beetle, all neonate individuals (body mass: 0.14mg) fed either leaves of the control line, line 69 (moderate MLX56-expressing line), or line 73 (high MLX56-expressing line) survived for 9 days, but the body size of the larvae fed leaves of line 69 was somewhat smaller than that of those fed leaves of the control, and the body size of larvae fed line 73 was much and obviously smaller than the size of those fed the leaves of the control line (Fig. 5A). The body weight of larvae fed line 69 for 9 days (12.77 ± 0.81 mg, average ± SD) was significantly but moderately smaller than those fed the control line (18.28 ± 0.98mg); that of the larvae fed leaves of line 73 (1.25 ± 0.12mg) was significantly smaller than those of both the larvae fed control leaves and the larvae fed leaves of line 69 (Fig. 5B) (p < 0.0001, *F* = 139.062). All larvae were 3^rd^ instar in the control and 69 lines, while only two larvae were 3^rd^ instar and the remaining 10 larvae were 2^nd^ instar in the 73 line. These results demonstrated that the expression of the MLX56 gene is also effective against the hadda beetle, and that the growth-inhibiting activities of MLX56-expressing lines are well correlated with the expression level of MLX56.

**Fig. 5.**
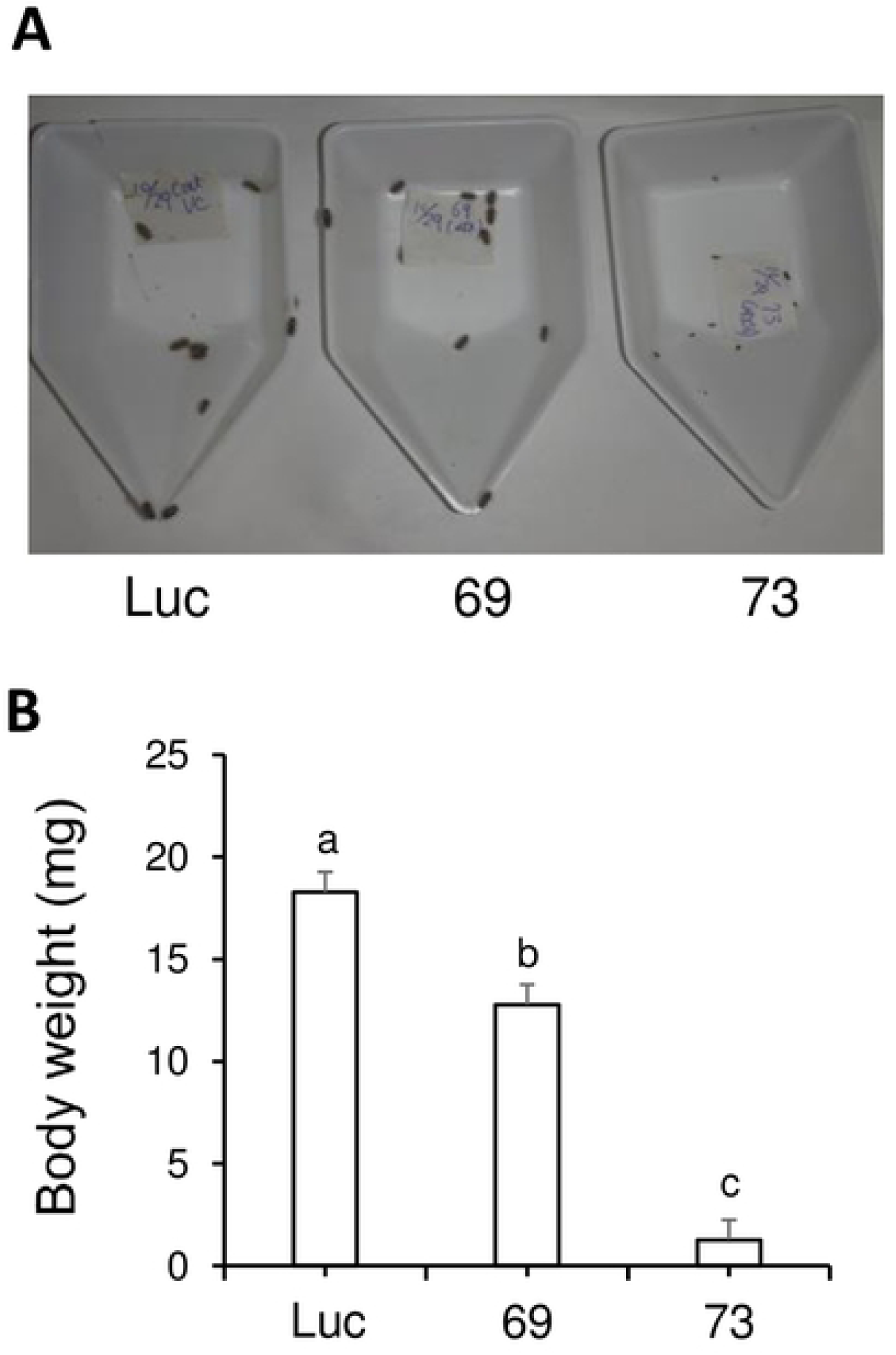
The effect of MLX56 on *H. vigintioctopunctata* larvae. Twelve newly hatched larvae (0.14 mg) were fed tomato leaves of either the control, 69, or 73 line for 9 days, and then, larval weights were then measured. Larval body size (A), and average body weights (B) on day 9. Error bars indicate SE (n = 12). Values not followed by the same letters were significantly different (Tukey’s test for multiple comparison, p < 0.0001, F = 139.1).

## Discussion

We produced transgenic tomato lines that overproduce the anti-herbivory protein MLX56. The transgenic tomato lines exhibited considerable tolerance against Lepidoptera larvae *S. litura*, as expected. The significant difference in the proportions of the larval instars on days 3 and 6 suggests that MLX56 serves to delay the development and/or reduce the survival rates during younger stages. Therefore, it is possible that with this effect, MLX56 protects crops from *S. litura* by preventing and/or delaying the emergence of older instar larvae that damage crops much more than younger instar ones. Further, these transgenic plants were remarkably resistant to *F. occidentalis* and *H. vigintioctopunctata*. The resistance of the line 73 tomato plant was so strong that the damage by thrips on the tomato line was negligible, and that the growth of hadda beetle was minimal. Therefore, MLX56 expression confers a resistance level to plants that is sufficiently strong to be practically utilized in agriculture like Bt toxin.

MLX56 was previously identified as an anti-herbivory defense protein of mulberry against lepidopteran herbivores [5, 11]. Although the molecular mechanism of MLX56 has not been fully determined, the protein is known to cause abnormal swelling of the PM, probably through its binding to chitin of the membrane, and the swollen PM is suggested to function as a barrier to the digestive processes of insects. The resistance level that MLX56 expression confers to plants is sufficiently strong to be practically utilized in agriculture like Bt toxin.

Surprisingly, ectopic expression of MLX56 also enhanced resistance against thrips, which have no PM [13]. At present, the mode of action and target of MLX56 in thrips are unclear, but chitin is everywhere in the body of all insects, including thrips, and there may be an additional/alternative chitin-containing targets of MLX56 in thrips, like the PM like structure in the midgut, cuticle in the foregut and hindgut, mouth parts, trachea, etc. The defensive mode of action of MLX56 in thrips should be examined in the future.

Constitutive high expression of the gene for MLX56 was achieved by using the expression vector pEl2Omega, which has 10-20 times higher activity than the CaMV 35S promoter, which is known to be strong in plants. The transgenic tomato line with higher expression of MLX56 (MLX56-73) exhibited higher resistance against thrips and beetles than the one with relatively less expression (MLX56-69), suggesting that a higher expression of MLX56 is required to provide high tolerance against insect pests. If the expression level is sufficiently high, the resistance level will be high enough to be effective against insect pests in practical agricultural use.

Various attempts have been made to produce transgenic crops with herbivore resistance by introducing plant-originated anti-herbivory genes; however, none are in practical commercial use at present. Meanwhile, the bacterial insecticidal gene BT has been successfully used to introduce herbivore resistance to crops worldwide. In particular, a considerable number of commercial cultivars of maize and soybean contain the BT gene, because it conveys potent resistance, and many types of BT gene have been reported. Despite its potent insecticidal activity, however, each type of BT has a relatively narrow action spectrum, and is effective only against limited types of pest insects. Therefore, a transgenic crop with one BT gene can resist only limited kinds of herbivores and multiple types of BT genes must be introduced to produce a transgenic crop with resistance against multiple herbivores. Further, the emergences of BT-resistant varieties of herbivore is frequently reported. Thus, potent anti-herbivory genes with a wide action spectrum and a different action mechanism than BT are needed to produce useful transgenic crops with resistance against various types of insect pests.

We consider that the gene for MLX56 could be a solution to the problem described above. Compared to GM plant production using Bt, the great advantage of producing pests resistant GM crops using MLX56 is that expressing a single MLX56 protein can confer sufficiently strong and broad resistance to important economical pests from various insect taxa such as thrips (Thysanoptera), ladybird beetles (Coleoptera), and cutworm (Lepidoptera) at the same time. Also, the performance of tomato as a crop does not seem to be compromised by the expression of MLX56. Therefore, introduction of the gene for MLX56 will be a promising and practical way to produce pest-resistant crops against a wide range of herbivores. Further, as the action mechanism of BT and MLX56 are different, simultaneous introduction of genes for BT and MLX56 may enlarge the action spectrum of the resistance and suppress occurrence of resistant varieties. We are currently introducing the gene for MLX56 under the control of an improved expression vector to evaluate the feasibility of MLX56 in different crops and herbivores.

## Acknowledgments

We thank Masumi Teruse, Chiaki Kimoto, Yuki Suzuki, Sachiko Gonokami, Kayoko Furukawa for technical assistance.

## Author Contributions

Conceptualization: Kotaro Konnno, Atsushi Mochizuki, Ichiro Mitsuhara.

Formal analysis: Mika Murata, Kotaro Konno, Ichiro Mochizuki.

Supervision: Ichiro Mitsuhara, Kotaro Konno.

Validation: Kotaro Konno, Naoya Wasano.

Writing – original draft: Mika Murata, Kotaro Konno, Ichiro Mitsuhara.

Writing – review & editing draft: Mika Murata, Naoya Wasano, Kotaro Konno, Ichiro Mitsuhara.

